# Contrasting topologies of synchronous and asynchronous functional brain networks

**DOI:** 10.1101/2024.07.23.604765

**Authors:** Clayton C. McIntyre, Mohsen Bahrami, Heather M. Shappell, Robert G. Lyday, Jeremie Fish, Erik M. Bollt, Paul J. Laurienti

## Abstract

We generated asynchronous functional networks (aFNs) using a novel method called optimal causation entropy (oCSE) and compared aFN topology to the correlation-based synchronous functional networks (sFNs) which are commonly used in network neuroscience studies. Functional magnetic resonance imaging (fMRI) time series from 212 participants of the National Consortium on Alcohol and NeuroDevelopment in Adolescence (NCANDA) study were used to generate aFNs and sFNs. As a demonstration of how aFNs and sFNs can be used in tandem, we used multivariate mixed effects models to determine whether age interacted with node efficiency to influence connection probabilities in the two networks.

After adjusting for differences in network density, aFNs had higher global efficiency but lower local efficiency than the sFNs. In the aFNs, nodes with the highest outgoing global efficiency tended to be in the brainstem and orbitofrontal cortex. aFN nodes with the highest incoming global efficiency tended to be members of the Default Mode Network (DMN) in sFNs. Age interacted with node global efficiency in aFNs and node local efficiency in sFNs to influence connection probability. We conclude that the sFN and aFN both offer information about functional brain connectivity which the other type of network does not.

## Introduction

There are many existing methods for studying functional brain connectivity using neuroimaging data (Smith et al., 2011), but the most commonly used methods are built upon the notion that near-synchronous (for simplicity, near-synchronous connections will be referred to as synchronous in this work) activation is indicative of functional connections. While traditional synchronous networks (Bullmore & Sporns, 2009) provide a useful backbone for understanding brain function, they are limited in that they do not capture the dynamic properties that functional brain networks are now known to possess (Hutchison et al., 2013; Lurie et al., 2020). Investigation of time-varying synchronous networks has become increasingly popular in recent years (Bassett & Sporns, 2017). Additionally, there are now a growing number of studies examining the brain’s ability to shift between metastable “states” of synchronous connectivity (Allen et al., 2014; Calhoun et al., 2014; Shappell et al., 2019; Shine et al., 2016; Zalesky et al., 2014). While these studies are excellent additions to our knowledge of brain function, they do not provide an explanation as to how the brain is able to self-direct shifts in functional connectivity.

We argue that understanding how synchronous networks change over time requires an understanding of asynchronous relationships between nodes. Specifically, we propose that changes in the brain’s synchronous relationships may be governed by asynchronous relationships, as has been shown recently in networks inspired by neural connectivity (Kumar et al., 2024). These asynchronous relationships can be modeled as a sparse directed network with the same nodes as the undirected synchronous network, but with edges distinct from those of the synchronous network both in distribution and meaning.

Previous neuroimaging studies have used Granger causality (Dhamala et al., 2008) and partial Granger causality (Guo et al., 2008) in attempts to demonstrate avenues of information flow through the brain. Transfer entropy-based approaches for studying information flow have also been applied in several previous studies (Maki-Marttunen et al., 2013; Palus et al., 2001; Rabinovich et al., 2012; Shovon et al., 2017). Granger causality and transfer entropy are both limited in that they cannot distinguish between direct versus indirect causal relationships. Partial Granger is extremely computationally involved and is not feasible for many researchers. One of the most popular approaches for understanding causal relationships in the brain is dynamic causal modeling (DCM)(Friston et al., 2003). However, due to the complexity of the method, DCM has been limited to brain networks with small numbers of brain regions and requires explicit hypotheses for selecting appropriate inferential procedures (Bahrami et al., 2023; Stephan et al., 2010). This limits its utility as a model of causality across the whole brain, especially when the underlying network topology is unknown.

Previous work outside of the network neuroscience literature has shown that causal networks can be inferred from time series data using optimal causation entropy (oCSE)(Sun & Bollt, 2014; Sun et al., 2015). This approach is intriguing as it utilizes concepts from information theory to determine direct causal relationships, distinct from indirect effects (Schreiber, 2000), while allowing for nonlinearity. In contrast to familiar synchrony-based functional networks, causation entropy-based directed networks are specifically conditioned to look beyond the synchronous relationships of nodes to determine causality. In this work, we refer to these synchrony-resistant causal networks as asynchronous functional networks (aFNs) to contrast them with the synchronous functional networks (sFNs) commonly studied in network analyses of neuroimaging data. Further, recent work has demonstrated a mathematical basis for how aFNs govern reorganization of sFNs (Kumar et al., 2024). It is not yet known whether the concept of asynchronous relationships which govern synchronous relationships applies to human brain function. Before this can be properly studied, it must be demonstrated that methods such as oCSE can be used to generate aFNs from human neuroimaging data, and these networks should add some biological meaning that is not already present in sFNs. The goal of the present work is to establish these foundations so that future work can explore whether aFNs do indeed act as governors of changes to synchronous relationships in the human brain.

In this work, we use resting state fMRI data from the NCANDA study (Brown et al., 2015) to generate sFNs and aFNs. Pairwise correlation was used to generate sFNs and oCSE was used to generate aFNs. Importantly, we have chosen to use oCSE to generate aFNs because unlike Granger causality or transfer entropy, it can distinguish direct associations from those with shared causal intermediaries. We compare the topology of the novel aFNs to the topology of the sFNs. Finally, we provide an example of how aFNs and sFNs can yield complementary information that would not be found with either network alone.

## Methods

### Data Collection

All data came from the National Consortium on Alcohol and Neurodevelopment in Adolescents (NCANDA), a longitudinal, multi-site neuroimaging study of adolescents aged 12-22 years at baseline (Brown et al., 2015). Only the baseline visits of participants from the University of California San Diego site were used for analyses. Because previous work using the NCANDA sample has reported scanner-related differences in fMRI data (Muller-Oehring et al., 2018), analyses were limited to data from one site to avoid confounding effects of different scanner models and site locations. The San Diego site was chosen for analyses as it had the most participants with baseline functional and structural MRI scans (n=212). The full NCANDA imaging parameters and protocols are described in previous publications (Muller-Oehring et al., 2018; Zhao et al., 2021). Briefly, the San Diego site used a 3T GE Discovery MR750 scanner with an 8-channel head coil. Subject motion was minimized using a head-strap with padding placed under the neck and along the sides of the participant’s head. High resolution (0.9375 mm × 0.9375 mm × 1.2 mm) T1-weighted structural scans were acquired using an Inversion Recovery-SPoiled Gradient Recalled (IR-SPGR) echo sequence (repetition time [TR] = 5.912 ms, echo time [TE] = 1.932 ms, 146 slices, acquisition time = 7 m 16 s). Resting state BOLD-weighted image sequences were acquired using a gradient-recalled echo-planar imaging sequence (TR = 2200 ms, TE=30 ms, 32 slices, acquisition time = 10 m 03 s). Subsequent sections of this work refer to TRs as specific time points in the functional image time series.

### Image Preprocessing

All image processing was completed in SPM 12 (https://www.fil.ion.ucl.ac.uk/spm/). Image processing began with unified segmentation (Ashburner & Friston, 2005) of T1-weighted structural images using standard six tissue priors while simultaneously warping images to Montreal Neurological Institute (MNI) standard space. The first 6 volumes (13.2 s) of the functional scan were removed to allow the signal to reach equilibrium. The remaining 269 volumes were corrected for slice-time differences, then realigned to the first volume. The previously defined warp to MNI space was then applied to the functional images. To further remove physiological noise and low frequency drift, functional data was filtered using a band-pass filter (0.009 – 0.08 Hz). Average whole brain gray matter, white matter, and cerebral spinal fluid mean signals, along with the six degrees of freedom motion parameters obtained from realignment were regressed from the functional data. Additionally, motion scrubbing (Power et al., 2014) was used to identify volumes with motion artifacts coupled with BOLD signal change, resulting in a binary vector of affected brain volumes. The resulting vector was then used as an additional motion regressor. This motion correction approach allowed us to retain the entirety of each participant’s functional scan, which was essential for creating our aFNs. The functional images were then parcellated into the functionally defined 268-region Shen atlas (Shen et al., 2013).

### Asynchronous Functional Network (aFN) Generation

Optimal causation entropy (oCSE)(Sun et al., 2015), an approach designed to identify sparse directed circuits between nodes in a network using causation entropy (Sun & Bollt, 2014), was used to generate aFNs. Causation entropy operates on the premise that through proper conditioning on carefully chosen subsets of nodes, the effects of intermediaries can be removed, and thus direct causal connections can be found. Using this conceptual framework, causation entropy from the set of nodes *J* to the set of nodes *I* conditioning on details of the set of nodes *K* is defined as

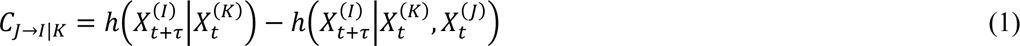

where h(*X*) is the Shannon entropy of *X*. To determine causation entropy of individual edges in a network, *J* = *j* and *I* = *i* and notation can be simplified to *C*_j→i|K_. That is, the causation entropy from node *j* to node *i* conditioning on another set of nodes, *K*. *K* represents the set of previously discovered (via a greedy algorithm, see Figure 1) neighbors of node *i*.

**Figure 1.**
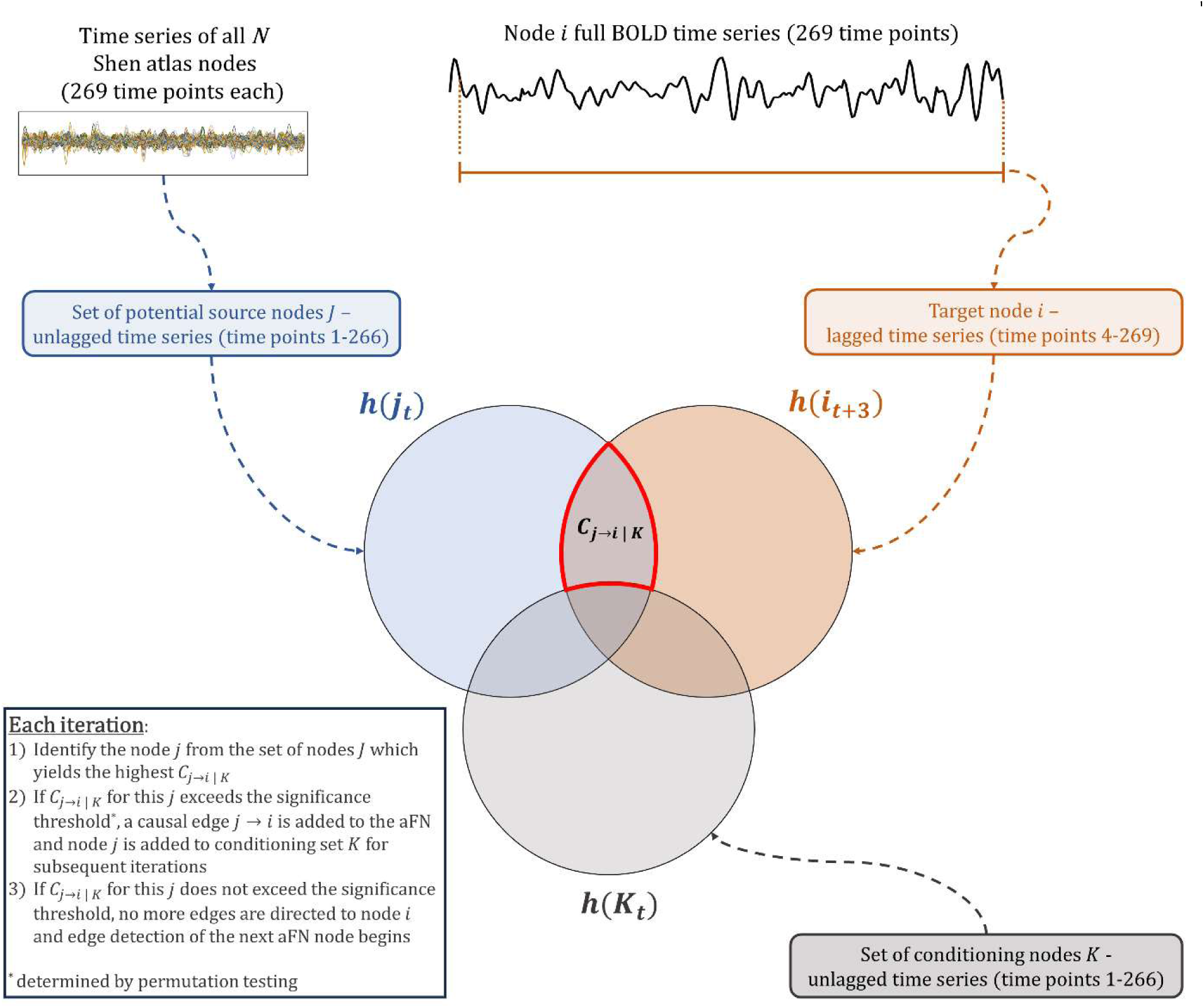
Schematic of oCSE detecting causal edges to node *i*. In oCSE-generated aFNs, causal edges to node *i* can come from any other node in the network, including node *i* itself. The target node’s time series is lagged (in this study, we used a uniform 3 time point lag) while all potentially causal edges are kept unlagged. oCSE identifies the unlagged node *j* with the highest conditional mutual information with the lagged target node *i.* For conditioning, a set of unlagged time series from nodes *K* is used. *K* starts as an empty vector and grows with each iteration until there are no more significant causal edges to node *i*.

This approach has previously been used to identify known structural connections using simulated fMRI time series data (Fish et al., 2021). For the first time, here we applied the causation entropy approach to real fMRI time series data to infer direct causal functional connections between nodes in the brain. Time series of each pairing of nodes were evaluated with one of the nodes having a time lag. For example, using a 1 TR time lag (τ = *t* + 1) in a 269 TR time series, the direct causal connection from node *j* to node *i* was determined by evaluating the causation entropy of *j*’s time series from TRs 1-268 and *i*’s time series from TRs 2-269. The time lag roles could then be reversed to determine whether *i* had a direct causal connection to *j*. This process was repeated for all possible pairings of nodes in the network, including self-connections (i.e., causation entropy of *j*’s time series from TRs 1-268 and *j*’s time series from TRs 2-269). After calculating the causation entropy of each possible edge, permutation testing (100,000 permutations) was applied to each edge to determine whether edges were statistically significant at α = 0.001. We chose this threshold because at this value, using the same time series data resulted in highly stable (i.e., similar) resulting networks. If desired, a higher threshold could be used, but this makes resulting networks more likely to have erroneous edges. Alternatively, a lower threshold could be used, but this would require more permutations and for the majority of researchers this would not be computationally feasible. After identifying whether edges between all pairs of nodes (in both directions) were significant, the result was a sparse, binary network in which only significant directed connections were present.

Multiple different TR time lags can be used to create causal networks. Previous work using time lagged embedding methods for a dynamic system found that the optimal time lag is the lag with the first minimum in mutual information between the original and lagged time series (Fraser & Swinney, 1986). For the current analyses the mutual information was computed for time lags of 1-5 TRs within the individual network nodes and the average for each lag was computed. The first minimum in the average mutual information was found at a lag of 3 TRs (τ = t+3, 6.6 seconds). This time lag also led to the fewest significant node self-connections in the aFN relative to other time lags tested. Therefore, aFNs built from a time offset of 3 TRs were used for all subsequent analyses in this study. For more details about different time offsets, see Supplementary Figure 1.

### Synchronous Functional Network (sFN) Generation

Pearson correlation was used on the time series of each possible pairing of two nodes resulting in a 268-node x 268-node correlation matrix. Functional brain network studies commonly apply proportional thresholds on correlation matrices, resulting in a binary adjacency matrix (Rubinov & Sporns, 2010). This approach sets each participant’s network to the same density (here density is used to describe the number of edges in a network divided by the number of edges in a fully connected network with the same number of nodes - thus, density is equal to 1-sparsity). For this study, sFNs were created solely to compare their topology to aFNs. For this purpose, proportional thresholding would be problematic because typical proportional thresholds used for sFN analyses (5-50% are not uncommon) are denser than aFNs (see Results). Thus, comparisons of topological features would be confounded by significant differences in network density, resulting in spurious results (van Wijk et al., 2010). Further, there is variability in aFN density across participants, so a standard proportional threshold across all sFNs may further confound comparisons between aFNs and sFNs. Therefore, we set each participant’s sFN to a personalized density level. Because the aFNs are very sparse, matching each participant’s sFN density to their aFN density would result in fragmented sFNs (reflected by a lower giant component – see Supplementary Table 1). Such fragmented networks would not be representative of the “small world” properties commonly reported for sFNs (Bassett & Bullmore, 2006; Sporns, 2013). Thus, each participant’s sFN was thresholded using the minimum number of edges needed to make the sFN giant component at least as large as the participant’s aFN giant component. The giant component is the largest connected component of the graph and its size is crucial for interpreting network efficiency measures (Latora & Marchiori, 2001) as a graph with disconnected nodes will tend to have much lower efficiency. The sFN for each participant was still denser than the aFN, so density was included in all statistical analyses to control for differences between the two types of networks. Supplemental analyses were also performed on sFNs using a range of proportional thresholds so that results could also be compared to existing literature (see Supplementary Table 1 and Supplementary Figure 3).

### Statistical Analyses – Comparing aFN and sFN Topology

The similarity of networks across participants was assessed using the Jaccard similarity coefficient. This is calculated with 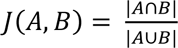 where *J*(*A,B*) is the Jaccard similarity coefficient of adjacency matrices *A* and *B*. That is, *J*(*A,B*) is an index value given by the shared edges between networks *A* and *B* divided by the total number of edges in *A* and *B*. The Jaccard similarity coefficient was calculated for every pairing of sFN adjacency matrices in the sample. These pairwise Jaccard values were then averaged resulting in one Jaccard similarity coefficient to represent the average similarity of all sFN adjacency matrices across the sample. This process was repeated for aFNs. The similarity of each participant’s sFN to their aFN was also assessed using Jaccard similarity. Jaccard values were then averaged across the sample to quantify the average similarity between a participant’s sFN and aFN with a single index value.

We used global and local efficiency (Latora & Marchiori, 2001) to quantify the ease with which information could be transferred between nodes in the sFNs and aFNs. Global efficiency of a node shows the average ease with which information can be transferred from a node to any other node in a network. Local efficiency of a node shows how easily information can flow between first neighbors of the node. Both values range from 0 (no edges) to 1 (edges with all other nodes) and can be averaged across all nodes to produce a network average level of global or local efficiency. Notably, while sFNs are undirected networks and global efficiency can be captured with a single value for each node, global efficiency can be calculated in aFN nodes in terms of either incoming or outgoing efficiency. While the same is technically true for local efficiency, the interpretation of directed local efficiency becomes challenging and potentially misleading, so we chose to calculate local efficiency of aFN nodes in terms of a full subgraph of first neighbors of both incoming and outgoing connections. Therefore, edge direction did not impact local efficiency values in the same way as global efficiency.

Efficiency values of sFNs and aFNs were calculated using the undirected and directed graph functions of the Brain Connectivity Toolbox (Rubinov & Sporns, 2010), respectively. To compare sFNs and aFNs in terms of network-wide efficiency (global and local), random intercept linear mixed effects models (to account for the repeated measures within a person) that controlled for the density of each participant’s sFN and aFN were used.

Pearson correlation was used to assess the relationship between the sample average efficiency values of nodes. For global efficiency in the aFN, values for incoming and outgoing efficiency were calculated separately and compared with each other as well as with sFN efficiency. Only one local efficiency value was computed for local efficiency for aFNs, and separately for the sFNs before using Pearson correlation on the pairing.

### Statistical Analyses – Comparing the Distribution of Highly Efficient Nodes

Differences in spatial distribution of efficiency were also assessed at the nodal level. Spatial mappings were created by overlaying the top 20% most globally efficient nodes from each participant’s network. This resulted in a brain map of nodes which are consistently among the most globally efficient nodes in a network. For aFN global efficiency, two mappings were created – one for incoming efficiency and one for outgoing efficiency. To statistically assess differences in the distribution of highly efficient nodes between sFNs and aFNs, we used pairwise permutations of network labels (i.e., sFN, aFN-outgoing, aFN-incoming) with 300,000 permutations according to a permutation testing framework (Simpson et al., 2013). This process was repeated for comparing distribution of high local efficiency in aFNs and sFNs, but with only one local efficiency value for both sFNs and aFNs.

### Statistical Analyses – Multivariate Mixed Regression Model

To determine whether sFN and aFN topology were related to age in our adolescent participants, a mixed-effects multivariate regression model desgined for brain network analyses (Bahrami et al., 2019) was used. This model allowed for determining *if and how* age was associated with sFN and aFN topology (in separate analyses) by providing statistical inference (i.e., p-values) and quantified relationships (i.e., positive/negative relationships). More specifically, this model assessed whether age interacted with global and local efficiency of nodes to influence the probability that the nodes are connected in either a sFN or aFN. Two separate models were run for aFNs – one assessed incoming global efficiency while the other assessed outgoing global efficiency. Both aFN models included local efficiency.

## Results

### Sample Characteristics

The UC San Diego site had 212 participants with structural and functional MRI scans at baseline. The sample was 48.6% female. The average age was 16.4 years with a range of 12.0-22.0 years. Most participants at this site identified as white (71.2%) and non-Hispanic (73.1%).

### Comparison of sFN and aFN Topology

Pairwise comparisons of sFNs across the sample resulted in an Jaccard similarity index of sFNs was 0.197 ± 0.027. For pairwise comparisons of aFNs across all participants, the average Jaccard similarity index was 0.018 ± 0.002. The average Jaccard similarity index between the sFN and aFN of an individual participant was 0.026 ± 0.005. This indicates that across the sample, edges in the sFNs were far more consistent than edges in the aFN. Additionally, the adjacency matrices for an individual person’s sFN and aFN tend to share more edges than the aFNs of two different people.

To visualize common connectivity patterns of each specific network type, the binarized adjacency matrix of all participants were summed and subsequently divided by the sample size to create sample-wide sFN matrices and aFN matrices (Figure 2). In these matrices, the value at each cell represents the percentage of participants that had the corresponding edge in their network. Notably, the consistency of edge distribution in sFNs was far higher than in aFNs. However, there were a few edges (especially self-connections) in the aFNs which were much more common across participants than other aFN edges. To make interpretation of the full aFN adjacency matrix possible, we set the upper limit of the color scale to 10%. The actual maximum value in aFNs was 50.5%, but the mean edge consistency was only 3.3%. The nodes are ordered along each axis to place nodes which typically are assigned to the same sFN subnetwork near each other. In the sFNs, there are clearly defined subnetworks of nodes with high interconnectivity that are consistently present across study participants, predominantly along the matrix diagonal. These boxes of synchronized nodes represent well-defined subnetworks, such as the Default Mode Network (DMN)(Andrews-Hanna, 2012; Buckner et al., 2008) and Central Executive Network (CEN)(Menon, 2011). In contrast to sFNs, the aFNs lack clear subnetworks along the diagonal of the matrix, but there are nodes which consistently have more outgoing connections than other nodes. These nodes generally belong to the basal ganglia and frontotemporal networks of the sFN. While there is some consistency in which nodes tend to have the most outgoing edges in aFNs, there is less of a noticeable pattern in terms of which nodes have the most incoming edges.

**Figure 2.**
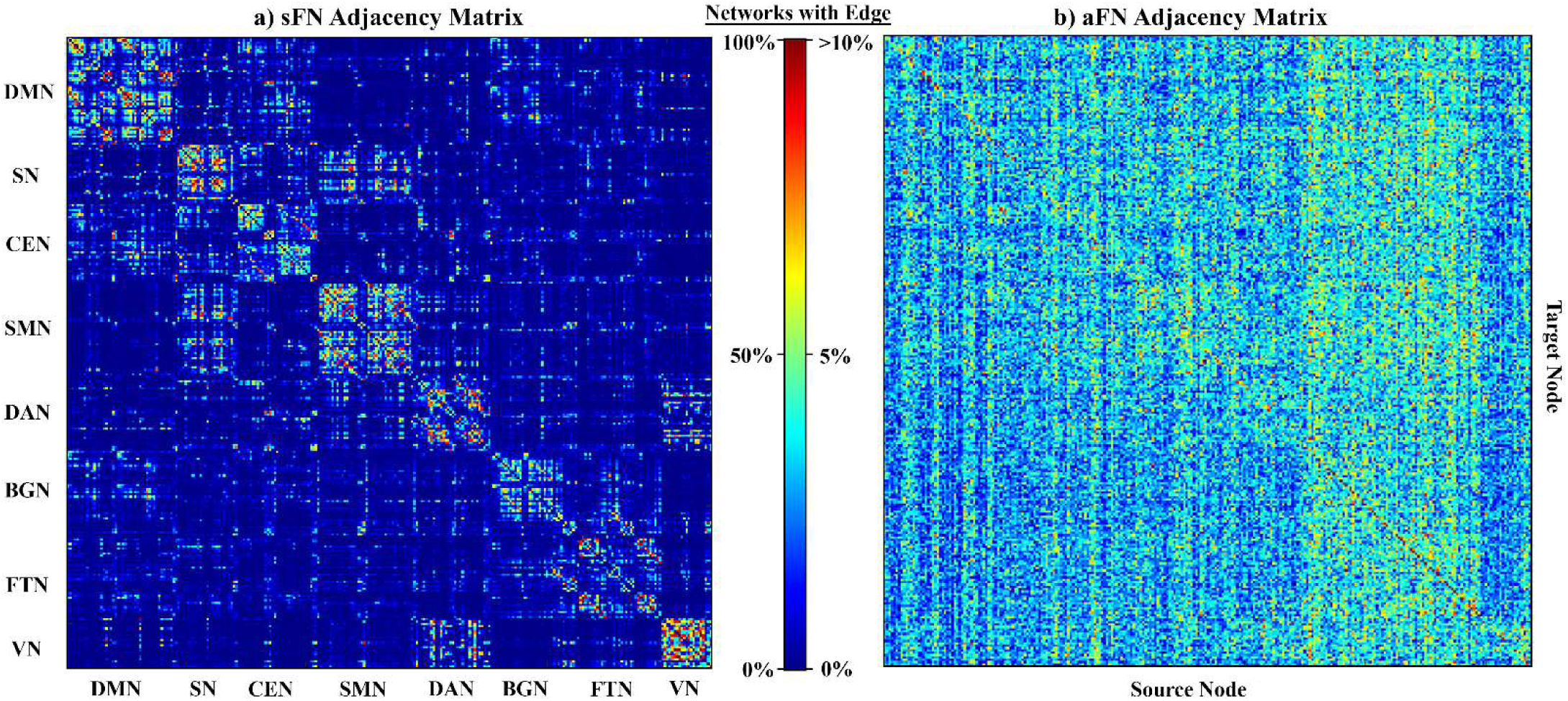
Sample-wide consistency of network edges. **a)**Sample-wide consistency of sFN adjacency matrices. The color scale shows the percentage of sFNs across the sample which include the edge. Distinct subnetworks can be discerned from the sFN adjacency matrix. (DMN = Default Mode Network, SN = Salience Network, CEN = Central Executive Network, SMN = Sensorimotor Network, DAN = Dorsal Attention Network, BGN = Basal Ganglia Network, FTN = Frontotemporal Network, VN = Visual Network). **b)** Sample-wide consistency of aFN adjacency matrices. Nodes are ordered identically to the matrix in a), but subnetworks are not labeled because they are not discernable from aFN adjacency matrices. There are noticeable vertical bands which indicate nodes with more outgoing connections than nodes in other subnetworks (these edges primarily originate from nodes that would be labeled part of the BGN and FTN in the sFN).

On average, the edge density of aFNs was 0.033 ± 0.005. The average edge density of the sFNs matched for the giant component size was 0.073 ± 0.024. Paired t-testing revealed that the density significantly differed between network types (p < 0.001, Figure 3a). Linear mixed effects modeling showed that after controlling for density differences, the mean global efficiency of an aFN was higher than the mean global efficiency of an sFN by 0.066 (p < 0.001, Figure 3b, Supplementary Table 2). The density-corrected analysis for local efficiency showed that the mean local efficiency of aFNs was lower than the mean local efficiency of sFNs by 0.545 (p < 0.001, Figure 3c, Supplementary Table 3).

**Figure 3.**
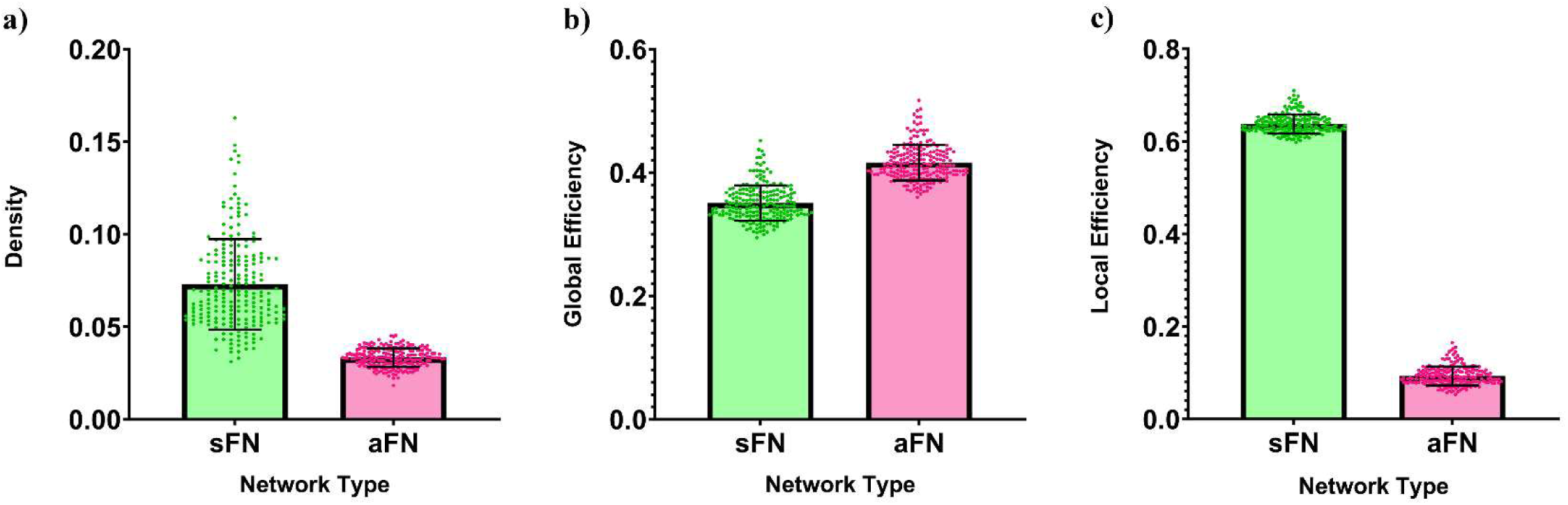
Comparison of density and network level efficiency between sFN and aFN. Individual points represent individual networks in the sample. All error bars represent standard deviation. Panel **a)** shows the density of sFNs and aFNs. Panel **b)** shows predicted global efficiency of sFNs and aFNs after adjusting for density differences between the networks. Values are calculated using parameter estimates of a random intercept linear mixed effects model with repeated measures. Each point represents a participant’s predicted sFN or aFN global efficiency with a network density equal to the average of the density of their personal sFN and aFN. Panel **c)** shows predicted local efficiency of sFNs and aFNs after adjusting for density as in (b), but using parameter estimates from the model with local efficiency instead of global efficiency.

### Distribution of Highly Efficient Nodes in the Brain

Figure 4 shows the spatial location of the nodes with high global efficiency (top 20%) across the sample for the sFN and for incoming and outgoing efficiency of the aFN. In the sFN, the nodes with high global efficiency were primarily in the frontal lobes, precuneus/posterior cingulate, and parts of the thalamus and cerebellum. aFN nodes with high outgoing global efficiency were mostly located in deeper brain structures such as the brainstem, cerebellum, basal ganglia, orbitofrontal cortex, and to a lesser extent the visual and sensorimotor cortex. aFN nodes with high incoming global efficiency were in the precuneus/posterior cingulate, posterior parietal lobes, and frontal lobes – all of which are hallmarks of the DMN in sFNs. Because this work is the first to use oCSE to generate aFNs, brain mappings were recreated using 1-year follow-up scans from the San Diego site as well as baseline scans of the Duke University site, both with very similar results to the baseline scans of the San Diego site (Supplementary Figures 3 and 4). Permutation testing revealed significant differences in the spatial distribution of nodes with high incoming and outgoing aFN global efficiency (p < 0.001), incoming aFN and sFN global efficiency (p < 0.001), and outgoing aFN and sFN global efficiency (p < 0.001). Pearson correlation showed that the average global efficiency of sFN nodes was negatively correlated with outgoing global efficiency of aFN nodes (r = −0.672, p < 0.001, Supplementary Figure 6a) but positively correlated with incoming global efficiency of aFN nodes (r = 0.161, p = 0.008, Supplementary Figure 6b). In aFN nodes, incoming and outgoing global efficiency was negatively correlated (r = −0.348, p < 0.001, Supplementary Figure 6c).

**Figure 4.**
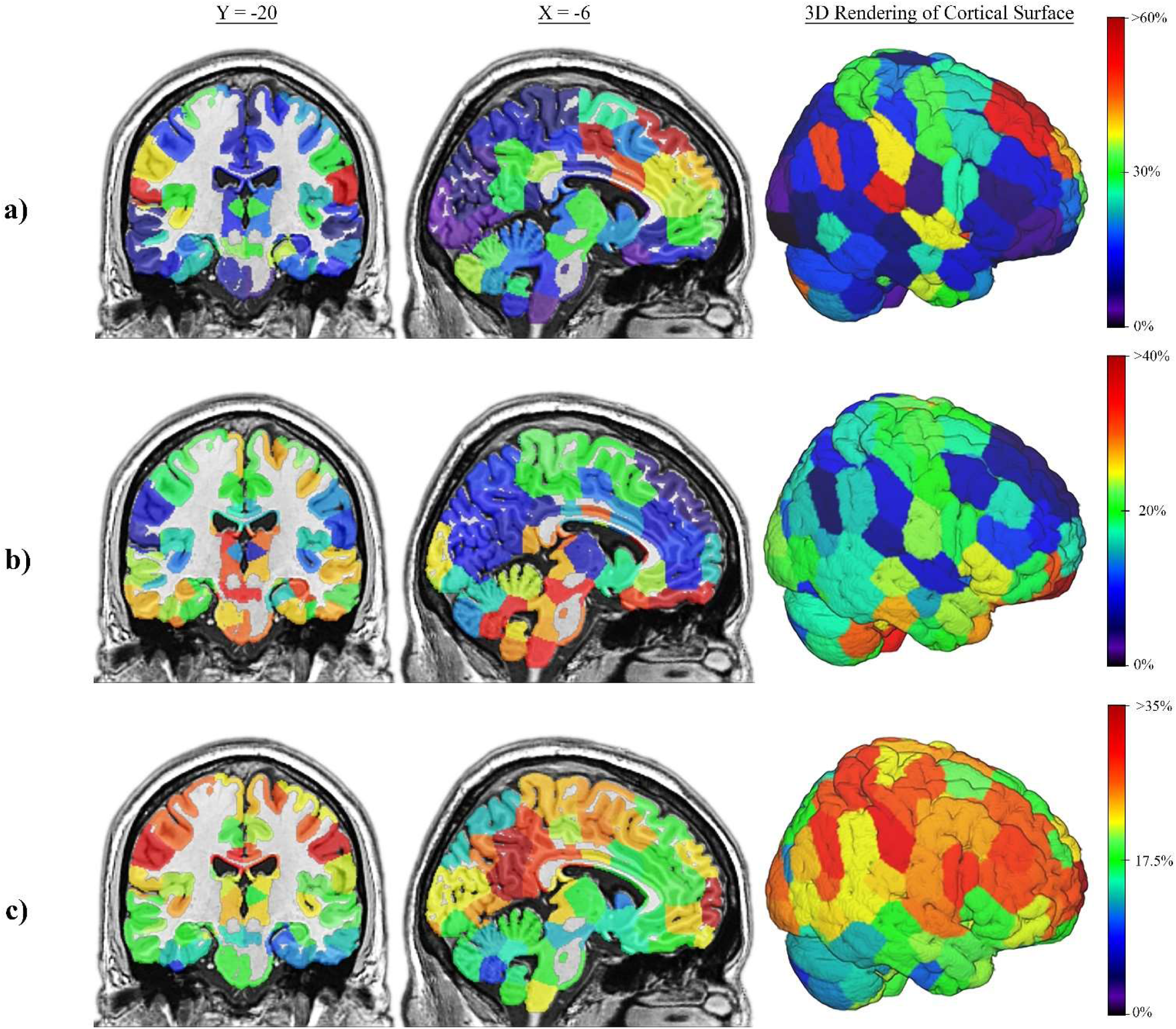
Spatial locations of high global efficiency nodes in the sFN and aFN. Brain maps depicting the location of nodes with high global efficiency across networks in the sample. Percentages on the color scale represent the percentage of networks in which a node appears in the top 20% of efficiency. Warmer colored nodes are more consistently among the most globally efficient nodes of a network relative to cooler colored nodes. MNI coordinates are shown above columns of each brain slice. Panel **a)** shows consistency of high global efficiency sFN nodes. Panel **b)** shows consistency of high outgoing global efficiency aFN nodes. Panel **c)** shows consistency of high incoming global efficiency aFN nodes.

Figure 5 shows the spatial location of the nodes with high local efficiency (top 20%) across the sample for the sFNs and aFNs. In the sFN, the nodes with high local efficiency were primarily in the visual cortex and parts of the parietal lobe. In the aFN, there was no noticeable pattern of where nodes with the highest local efficiency were. Figure 5 shows that the aFNs have much less consistency in the spatial distribution of high local efficiency relative to sFNs. The spatial distribution of high local efficiency nodes differed significantly between aFNs and sFNs (p < 0.001) as determined by permutation testing. Pearson correlation showed that the average local efficiency of sFN nodes and aFN nodes were not significantly related (r = −0.037, p = 0.546, Supplementary Figure 7).

**Figure 5.**
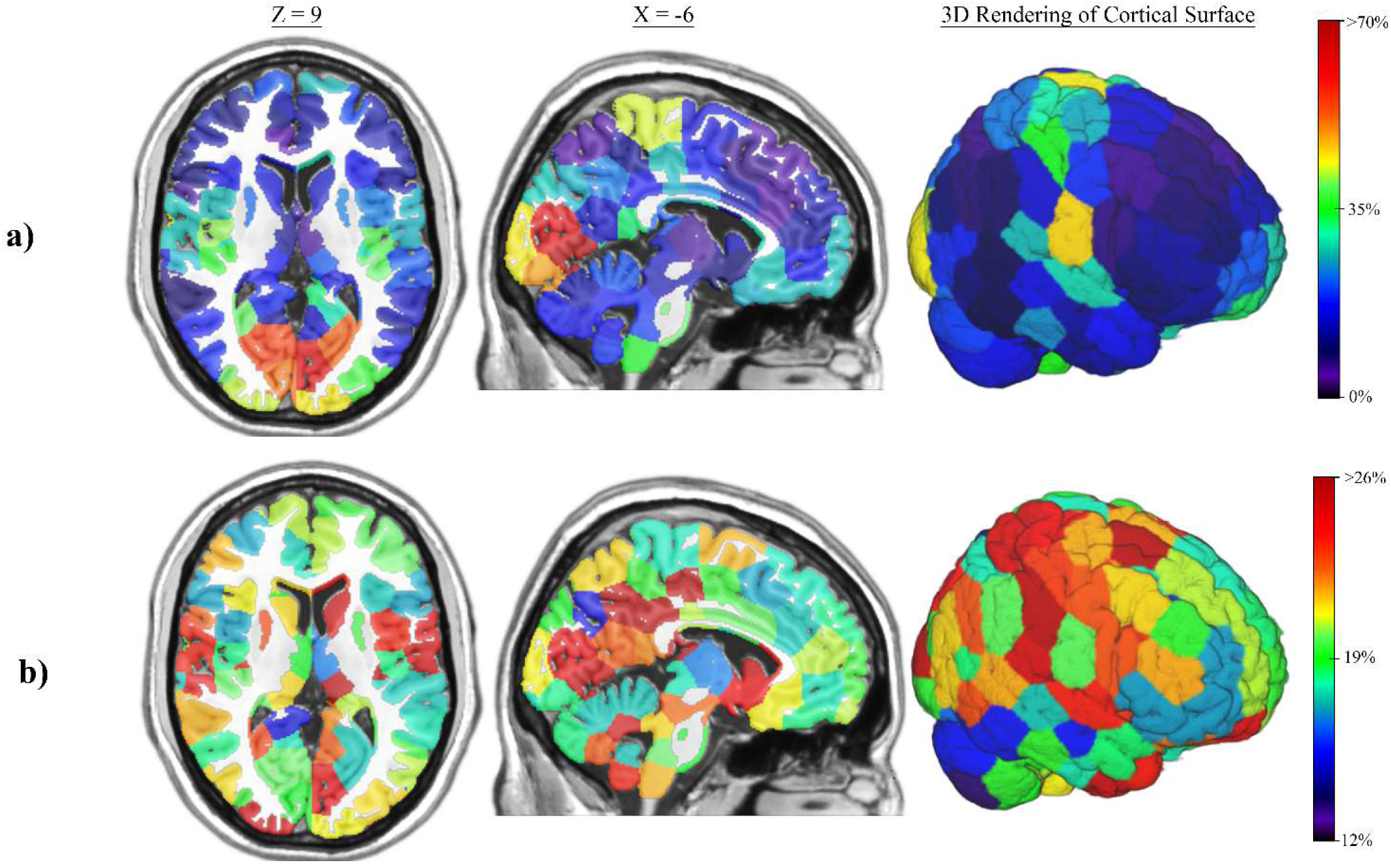
Spatial locations of high local efficiency nodes in the sFN and aFN. Brain maps depicting the location of nodes with high local efficiency across networks in the sample. Percentages on the color scale represent the percentage of networks in which a node appears in the top 20% of efficiency. Warmer colored nodes are more consistently among the most locally efficient nodes of a network relative to cooler colored nodes. MNI coordinates are shown above columns of each brain slice. Panel **a)** shows consistency of high local efficiency sFN nodes. Panel **b)** shows consistency of high local efficiency aFN nodes.

### Associations Between Network Topology and Age

To examine the relationship between brain network topology and age, we used a multivariate mixed effects regression analysis developed for brain network analyses (Bahrami et al., 2019). Our results in the aFNs revealed that as the outgoing global efficiency of two nodes increases, the probability of a connection between the nodes also increases. Age exhibited a significant interaction in this model (*β* = −0.016, SE = 0.006, *p* = 0.004, full results in Supplementary Table 4a) such that the relationship between outgoing global efficiency and connection probability was stronger for younger participants than for older participants (Figure 6a). A similar result was found for incoming global efficiency in the aFN. That is, as the incoming global efficiency of two nodes increases, the probability of a connection between the nodes also increases. Age exhibited an interaction in this model (β = −0.014, SE = 0.006, p = 0.028, full results in Supplementary Table 4b) such that the relationship between incoming global efficiency and connection probability was stronger for younger participants than for older participants (Figure 6b). This result was validated in the same study site at one year follow up and in the baseline visits of the Duke University NCANDA site. While sample sizes were slightly smaller, results at these sites/visits were similar to the baseline results for the San Diego site (Supplementary Tables 5 and 6). There was no statistically significant relationship between age and global efficiency in the sFNs (β = −0.008, SE = 0.008, *p* = 0.313, full results in Supplementary Table 7).

**Figure 6.**
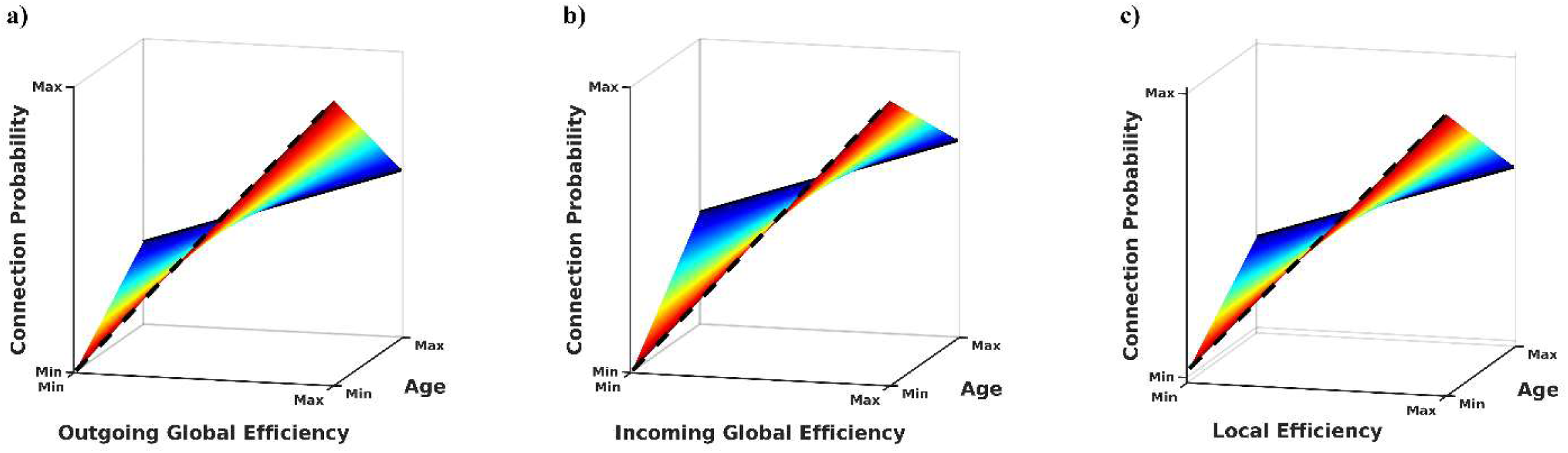
Age-nodal efficiency interaction and connection probability. Panel **a)** shows the interaction between age and outgoing global efficiency of aFN nodes. As the outgoing global efficiency of two nodes increases in the aFN, the probability of a connection between the edges also increases, but this relationship is weaker (shallower slope) in older participants. Panel **b)** shows the interaction between age and incoming global efficiency of aFN nodes. As the incoming global efficiency of two nodes increases in the aFN, the probability of a connection between the edges also increases, but this relationship is weaker (shallower slope) in older participants. Panel **c)** shows the interaction between age and local efficiency of sFN nodes. As the local efficiency of two nodes increases in the sFN, the probability of a connection between the edges also increases, but this relationship is weaker (shallower slope) in older participants.

The same multivariate mixed effects model showed that the probability of a connection between two nodes in the sFN increased as the local efficiency increased. Age exhibited a significant interaction in this model (*β* = −0.020, SE = 0.007, *p* = 0.008, full results in Supplementary Table 6) such that the relationship between local efficiency and connection probability was stronger for younger participants than for older participants (Figure 6c). There was no statistically significant relationship between age and local efficiency in the aFNs (*β* = 0.003, SE = 0.004, *p* = 0.426, full results in Supplementary Table 4).

## Discussion

We have introduced a new approach for identifying sparse directed networks from time series data to the network neuroscience literature. We have compared the topology of networks generated with this approach to the topology of synchronous correlation-based networks, and we have demonstrated that aFNs capture unique network features which are not present in sFNs. While the methods and results of this work focus on one method for generating aFNs (i.e., optimal causation entropy) and one method for generating sFNs (i.e., pairwise Pearson correlation), we do not claim that these are the only valid approaches for creating these networks. We chose to use oCSE because it is a powerful, nonlinear method for detecting aFNs. We used correlation to generate sFNs because it is a dominant methodology in existing neuroscience literature. We argue that by considering both the synchronous connections between brain regions using sFNs and the asynchronous, causal connections between brain regions using aFNs, we are able to create a more meaningful picture of brain function than from either method alone, regardless of the specific method used to generate aFNs and sFNs. We believe that this combined approach may be more useful than focusing solely on synchronous relationships, as is currently the norm. We elaborate on the results presented in this work below.

Nodes which encompassed the brainstem nuclei tended to have high outgoing global efficiency. These nuclei are responsible for widespread neuromodulation (Benarroch, 2018; Hornung, 2003; Morales & Margolis, 2017; Venkatraman et al., 2017), and are likely key players in regulating neural dynamics. The visual cortex and parts of the somatosensory cortex also tended to have high outgoing efficiency. These findings may suggest that visual and bodily information are influential over many neural processes. Other nodes with high outgoing global efficiency include those in the orbitofrontal cortex and basal ganglia, both of which are thought to be crucial for reward prediction and behavioral automaticity and are highly relevant to informing motivated behavior and decision-making (Brown & Marsden, 1998; Graybiel, 2000; Ring & Serra-Mestres, 2002; Rolls, 2004).

Nodes with high incoming global efficiency were largely concentrated in the precuneus/posterior cingulate, dorsal medial prefrontal cortex, and lateral parietal cortex. All of these regions are core members of the DMN in sFNs. This subnetwork is thought to be involved in a wide range of cognitive processes, including self-referential thought (Andrews-Hanna, 2012). Additionally, several other nodes in the frontal and parietal lobes tended to have high incoming global efficiency. These regions are commonly associated with subnetworks involved in higher level cognitive processing, such as the Central Executive Network (CEN)(Menon, 2011). Taken together, these results may be indicative of a pattern in aFNs: sources of information flow tend to be key players in neuromodulation and early sensory processing, while targets of information flow tend to be crucial members of a higher-level cognitive processing “backbone” (Andrews-Hanna et al., 2010).

Our analysis of the age interaction on the relationship between network efficiency and connection probability had two motivations: 1) to demonstrate an approach for statistically interpreting the topology of aFNs, and 2) to demonstrate that aFN topology is significantly related to a biological variable that one might reasonably expect to influence brain function. We showed that age interacted with global efficiency and connection probability in aFNs, but not sFNs. This makes sense given that aFNs are characterized by their many “long-range” connections, which are reflected by their high global efficiency. A meaningful difference in aFN topology may therefore be easier to identify using a metric like global efficiency. We also showed that age exhibits an interaction with local efficiency and connection probability in sFNs - but not aFNs. This also makes sense given that sFNs are characterized by high clustering (and, similarly, high local efficiency), which supports functional specialization of subnetworks (Bassett & Bullmore, 2006). While it is possible that local efficiency may have an important role in aFNs, such a role is not immediately apparent.

Because this work is the first to apply oCSE to neuroimaging data, we conclude our discussion of results with notes on interpretation of oCSE-generated aFNs as sparse, directly causal networks. While oCSE is designed to identify directly causal relationships, the causal relationships shown in aFNs are not necessarily indicative of physiological/structural connections between connected nodes. Rather, edges in aFNs demonstrate that present activity in the source node directly influences the probability of observing future activity in the target node. Identifying neuromechanistic underpinnings of causal relationships is beyond the scope of this method. Additionally, we note that node pairings which exhibit low synchrony are often considered “weak” connections and disregarded in network analyses. However, even connections which appear weak from a synchrony-centric perspective can be crucial to the behavior of complex networks (Granovetter, 1973), including neural systems (Pessoa, 2014). aFNs may provide a new approach for identifying relationships between nodes which are not highly synchronized, but which are nonetheless functionally connected across time in a manner that is crucial for network dynamics.

There are limitations to the proposed approach of studying aFNs and sFNs together with currently available methods. First, the present study suggests that at the individual edge level, aFNs vary substantially across people. However, brain mappings (Figure 4) and multivariate mixed model results (Figure 6) suggest that at the network level, there are common characteristics across people. Therefore, the optimal approach for analyzing aFNs is likely through graph-theoretical analyses rather than comparison of individual edges. Another challenge to aFN generation and interpretation is the complexity of interpreting hemodynamic signal through TR lags, as has been noted in other work (Smith et al., 2011). Further, the brain is known to work at many different time scales (Buzsáki, 2006; Hasson et al., 2008); therefore, our application of a uniform 3 TR lag to generate aFNs may result in overlooking meaningful asynchronous connections that are detectable only with different time lags. The optimal time lag to apply likely depends on the specific research question and may vary between brains. We leave further exploration and discussion of this subject to future work. Finally, oCSE was designed for stationary Markov processes. The brain’s functional connectivity is not stationary. Therefore, it may not be accurate to assume that one aFN can sufficiently model the underlying causal relationships present in the brain across an entire fMRI scan. However, just as synchronous relationships across the whole brain were initially studied using static networks generated from correlation of the time series from entire brain scans (Bullmore & Sporns, 2009), study of aFNs needs to start somewhere.

We intended for this work to serve as a starting point for future efforts to explore these networks as well as to refine methodology associated with them. Ultimately, we hope that studying aFNs in addition to sFNs will allow researchers to view functional brain connectivity from a more holistic perspective with consideration for asynchronous as well as synchronous relationships.

## Acknowledgements

All data presented in this work was collected from baseline visits of the UC San Diego site of the NCANDA study. NCANDA is supported by NIH Grants AA021697, AA021697-04S1, AA021695, AA021692, AA021696, AA021681, AA021690, and AA021691. The NCANDA dataset is publicly available upon request (see http://www.ncanda.org/index.php).

