## Supplement for "Contrasting topologies of synchronous and asynchronous functional brain networks"

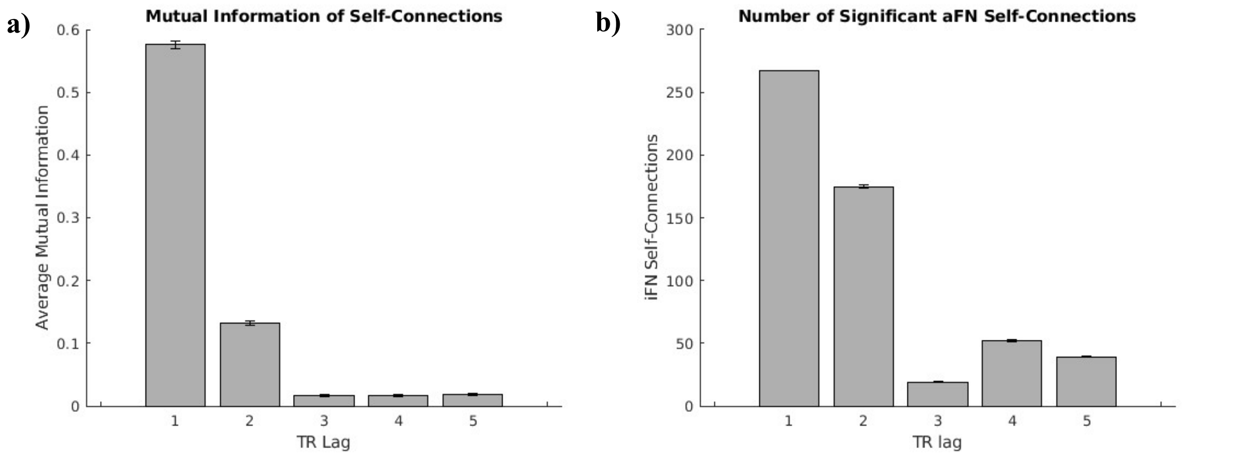


**Supplementary Figure 1. Time lag mutual information within the same node**

Error bars represent standard error. At some lags, standard error is too small to visually render in the figure.

(a) Average mutual information in the time series of a node at time *t* and itself at time *t+lag*.
(b) Total number of significant aFN self-connections with TR lags of 1-5 applied. A 3 TR lag produced the first minimum in number of self-connections in the aFN.

| Network Generation Approach | Density | Local Efficiency | Global Efficiency | Modularity | Giant Component |
|---|---|---|---|---|---|
| **aFN - TR Lag 3** | 0.033 (0.005) | 0.063 (0.016) | 0.374 (0.024) | 0.214 (0.020) | 266.8 (2.3) |
| **sFN - Matching aFN Giant Component** | 0.073 (0.024) | 0.668 (0.042) | 0.392 (0.057) | 0.581 (0.066) | 266.8 (2.3) |
| **sFN - Matching aFN Density** | 0.033 (0.005) | 0.567 (0.040) | 0.261 (0.040) | 0.673 (0.040) | 252.7 (11.7) |
| **sFN - Proportional Threshold (5%)** | 0.050 | 0.640 (0.020) | 0.342 (0.019) | 0.608 (0.031) | 263.6 (3.8) |
| **sFN - Proportional Threshold (10%)** | 0.100 | 0.710 (0.016) | 0.465 (0.012) | 0.492 (0.030) | 267.7 (0.7) |
| **sFN - Proportional Threshold (25%)** | 0.250 | 0.753 (0.013) | 0.619 (0.004) | 0.331 (0.029) | 268 (0) |

**Supplementary Table 1. aFN and sFN Network Metrics**

Values are presented as mean (standard deviation). Row 1 shows sample average aFN topology metrics. Row 2 shows the topology metrics used for analyses throughout the paper. In this approach, each participant's sFN density is unique to the individual such that the giant component of the sFN is minimized while satisfying the condition that it is at least as large as the giant component of the participant's aFN. With this approach, density tends to be roughly twice as high in sFNs as aFNs, but because the giant component is the same between the two networks it is more straightforward to compare the topology of the two. Row 3 shows sample average sFN topology metrics when each participant's sFN has its density matched to the participant's aFN density. Rows 4, 5, and 6 show sFN topology metrics when proportional thresholds of 5%, 10%, and 25% were applied, respectively.

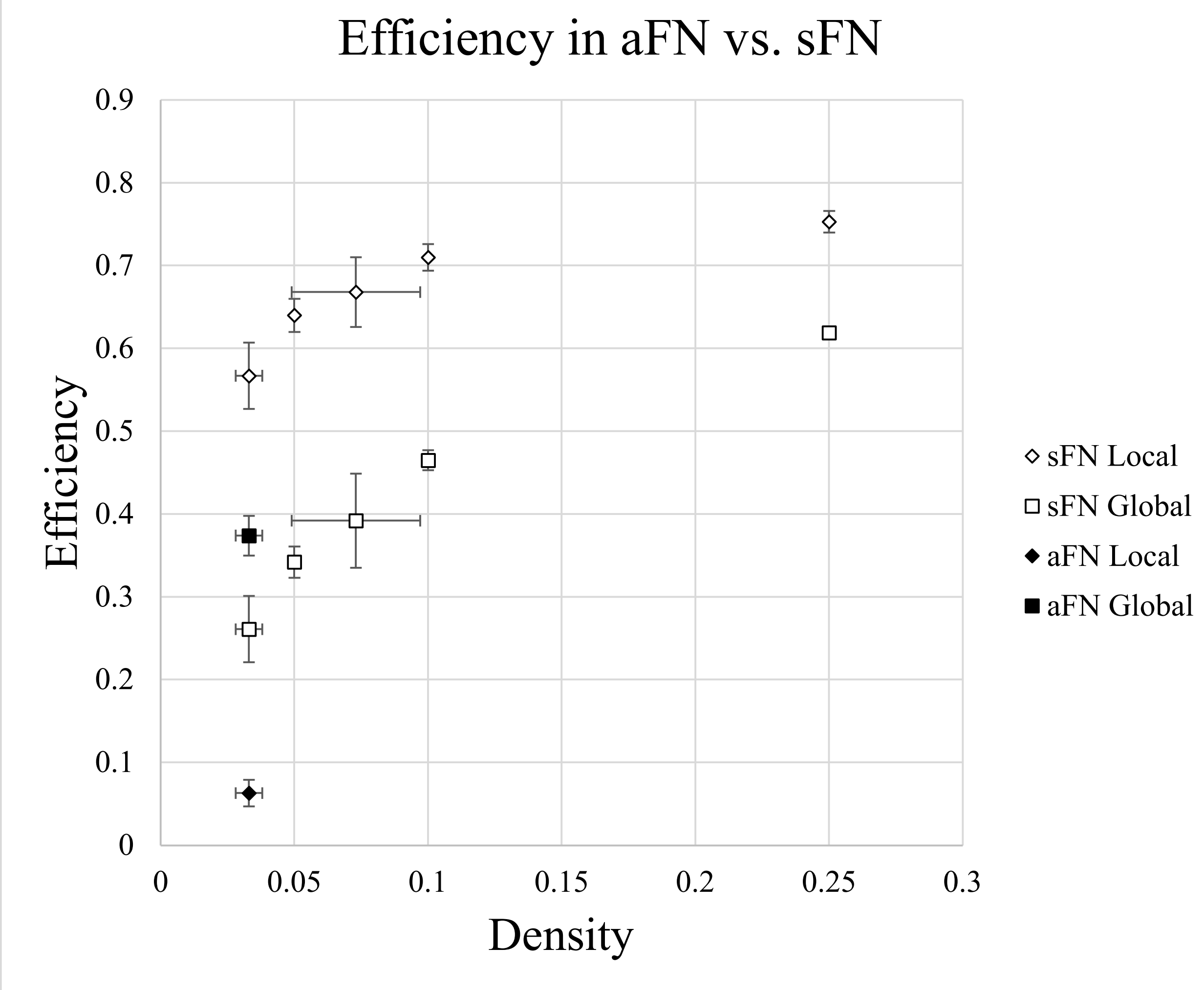


**<u>Supplementary Figure 2. Efficiency in aFN vs. sFN with Different Densities</u>**
Local and global efficiency of aFNs and sFNs of different densities. aFNs are created unique to each individual with density that cannot be readily modified. In our sample, the average density of aFNs was 0.033. Relative to sFNs with the same density as the participant's aFNs, the aFN is more globally efficient but less locally efficient. Both global and local efficiency increase in sFNs as the threshold for binary edges is raised and sFNs become denser. In some data points, standard deviation is too small to visibly render error bars.

| Fixed Effects | | | | | |
|---|---|---|---|---|---|
| **Effect** | **Estimate** | **Standard Error** | **DF** | **t Value** | ***p*** |
| **Intercept** | 0.303 | 0.002 | 420.931 | 163.22 | <0.001 |
| **Network Type** | -0.066 | 0.002 | 301.455 | -31.80 | <0.001 |
| **Density** | 2.122 | 0.043 | 359.941 | 49.55 | <0.001 |

**Supplementary Table 2. Linear Mixed Effects Model – Global Efficiency**

Linear mixed effects model with repeated measures was used to predict global efficiency of a network while controlling for network type (sFN = 1, aFN = 0) and network density. The estimate of "Network Type" is the predicted difference in global efficiency in sFNs relative to aFNs after accounting for density effects.

| Fixed Effects | | | | | |
|---|---|---|---|---|---|
| **Effect** | **Estimate** | **Standard Error** | **DF** | **t Value** | ***p*** |
| **Intercept** | 0.013 | 0.002 | 421 | 6.444 | <0.001 |
| **Network Type** | 0.545 | 0.003 | 421 | 212.995 | <0.001 |
| **Density** | 1.508 | 0.048 | 421 | 31.291 | <0.001 |

**Supplementary Table 3. Linear Mixed Effects Model – Local Efficiency**

Linear mixed effects model with repeated measures was used to predict local efficiency of a network while controlling for network type (sFN = 1, aFN = 0) and network density. The estimate of "Network Type" is the predicted difference in local efficiency in sFNs relative to aFNs after accounting for density effects.

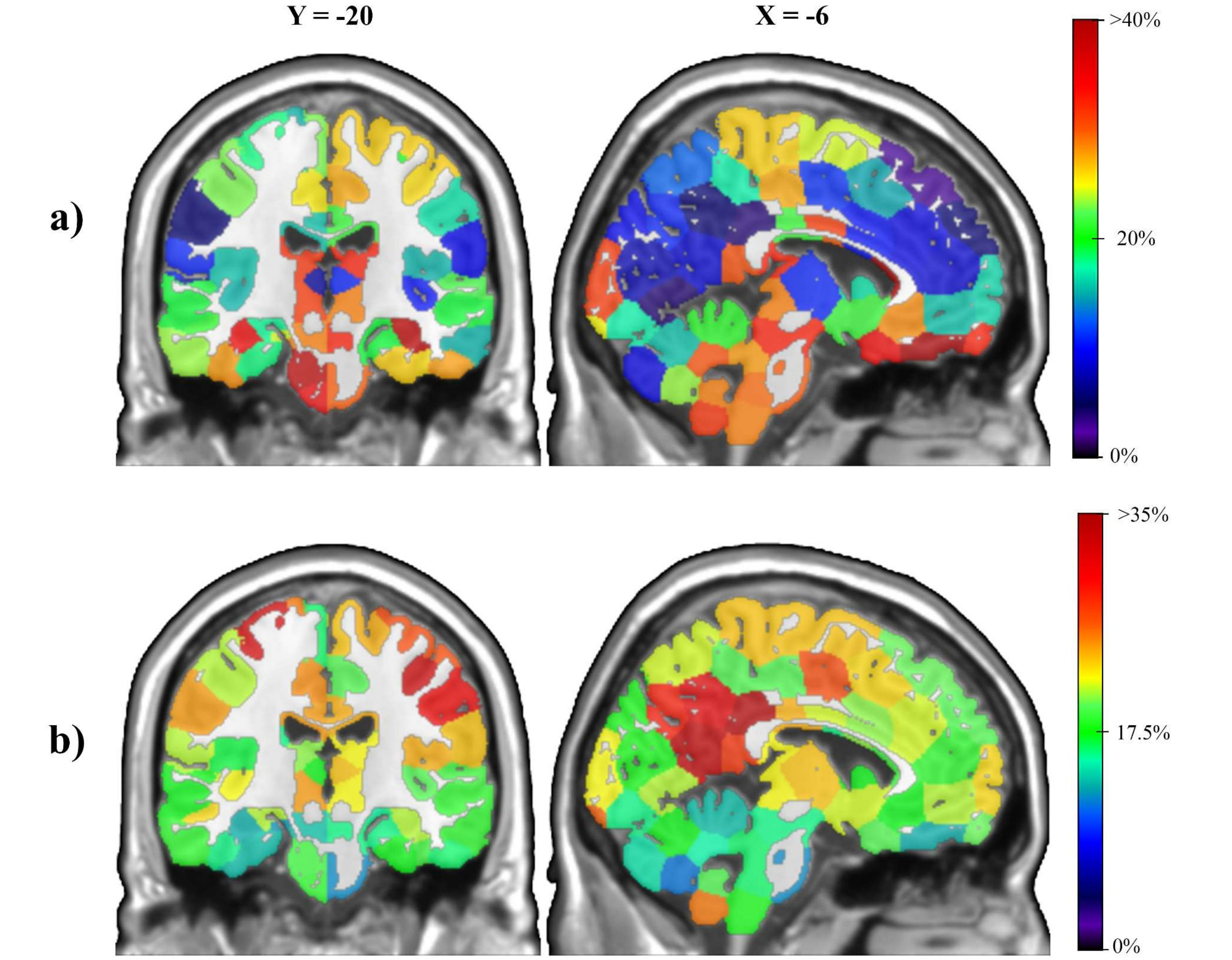


**Supplementary Figure 3. Spatial locations of high global efficiency nodes in San Diego site at one year follow-up**

Brain maps depict the location of nodes with consistently high global efficiency across the sample. Percentages on the color scale represent the percentage of networks in which a node appears in the top 20% of efficiency. Warmer colored nodes are more consistently among the most globally efficient nodes of a network relative to cooler colored nodes. MNI coordinates are shown above columns of each brain slice. Panel **a)** shows consistency of high outgoing global efficiency. Panel **b)** shows consistency of high incoming global efficiency. Spatial mappings of the 197 participants from the San Diego site at one year follow-up are very similar to mappings from the baseline visit (Figure 4).

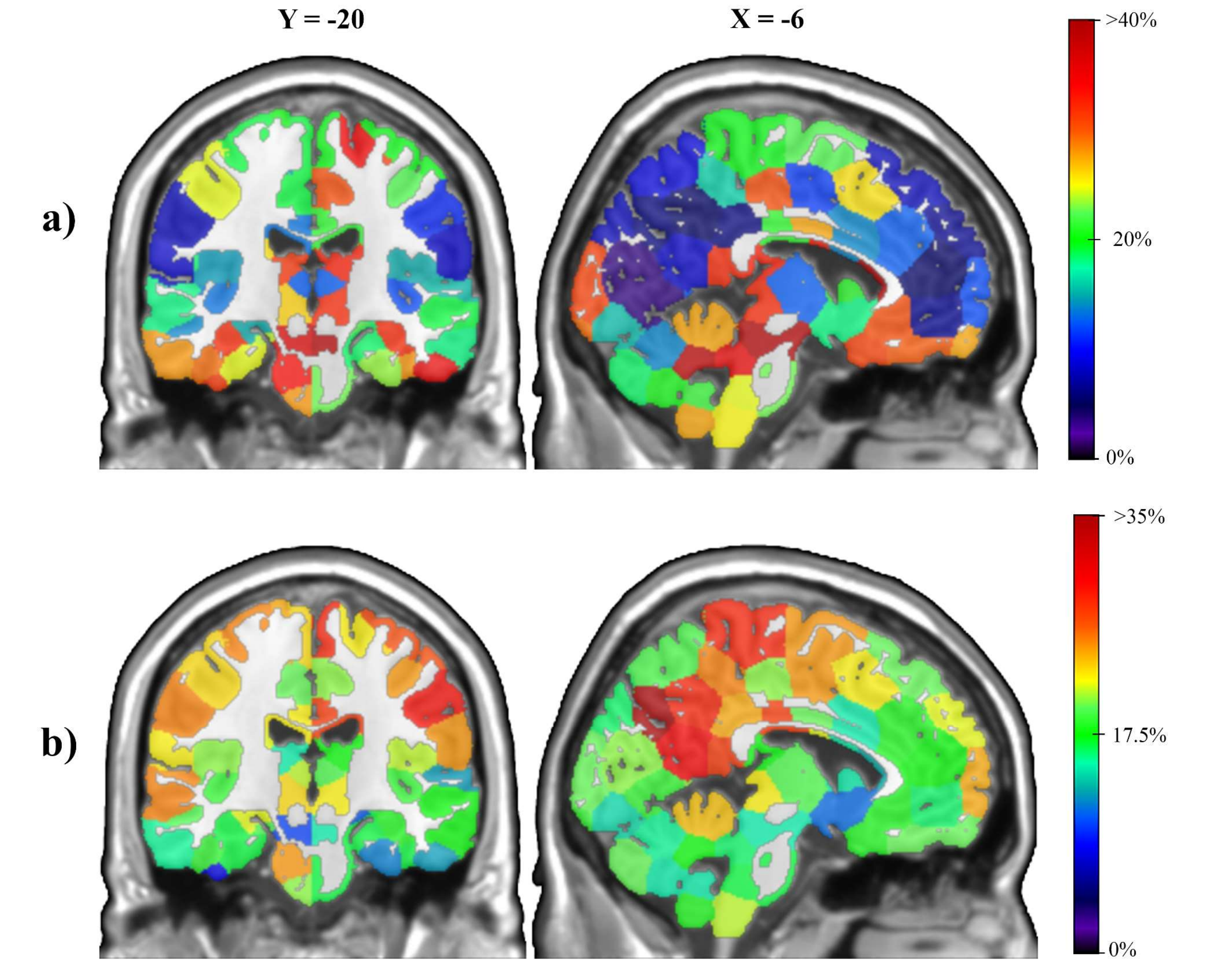


**<u>Supplementary Figure 4. Spatial locations of high global efficiency nodes in Duke site at baseline</u>**

Brain maps depict the location of nodes with consistently high global efficiency across the sample. Percentages on the color scale represent the percentage of networks in which a node appears in the top 20% of efficiency. Warmer colored nodes are more consistently among the most globally efficient nodes of a network relative to cooler colored nodes. MNI coordinates are shown above columns of each brain slice. Panel **a)** shows consistency of high outgoing global efficiency. Panel **b)** shows consistency of high incoming global efficiency. Spatial mappings of the 166 participants from the Duke site at baseline are very similar to mappings from the baseline visit of the San Diego site (Figure 4).

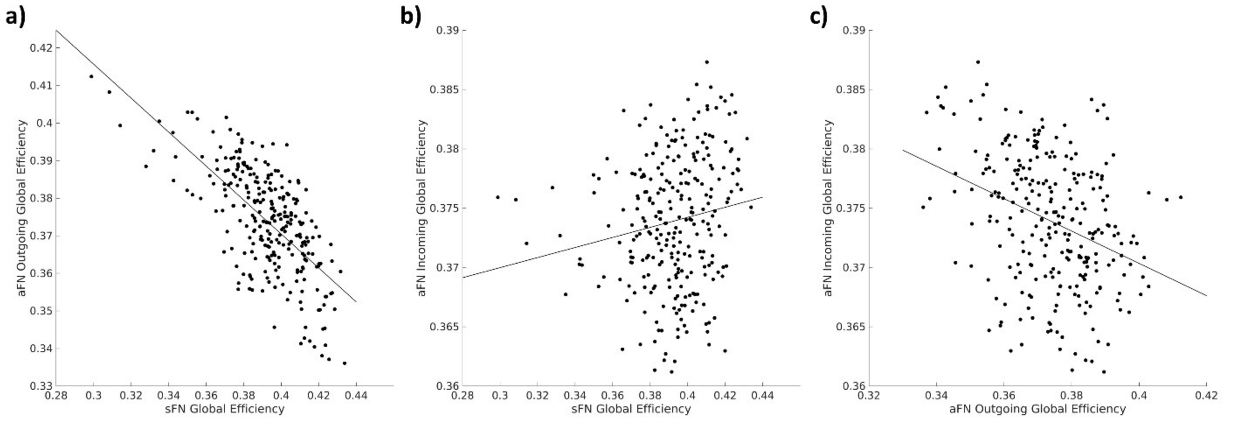


**<u>Supplementary Figure 5. Correlation of Node Global Efficiency in sFNs and aFNs</u>**
Scatterplot of the average global efficiency values in each node of the aFN and sFN. Each point in the figure represents one of the 268 nodes from the Shen atlas.

a) Comparison of sFN global efficiency to aFN outgoing global efficiency. Node values were significantly negatively correlated ($r = -0.672$, $p < 0.001$).
b) Comparison of sFN global efficiency to aFN incoming global efficiency. Node values were significantly positively correlated ($r = 0.161$. $p = 0.008$).
c) Comparison of aFN outgoing global efficiency to aFN incoming global efficiency. Node values were significantly negatively correlated ($r = -0.348$, $p < 0.001$).

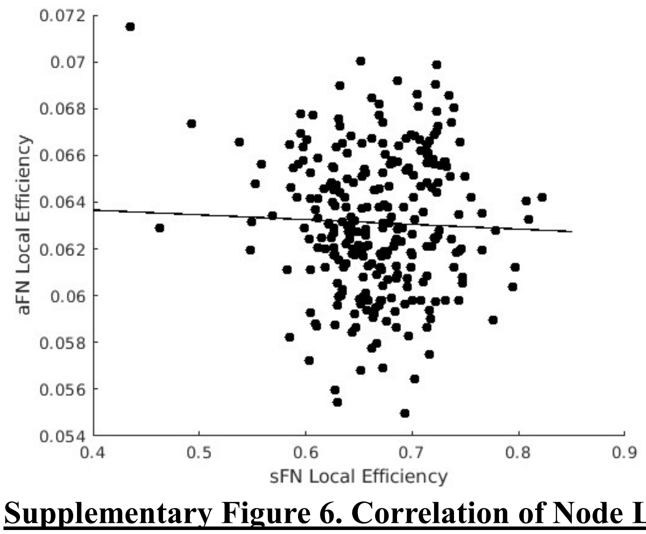


**<u>Supplementary Figure 6. Correlation of Node Local Efficiency in sFNs and aFNs</u>**
Scatterplot of average local efficiency values in each node of the aFN and sFN. Each point in the figure represents one of the 268 nodes from the Shen atlas. Local efficiency in the aFN and sFN were not significantly correlated ($r = -0.037$, $p = 0.546$).

| 4a. Solutions for Fixed Effects – aFN Model 1 (Outgoing Connections) | | | | | |
|---|---|---|---|---|---|
| **Effect** | **Estimate** | **Standard Error** | **DF** | **t Value** | **p** |
| **Intercept** | -3.268 | 0.009 | 152E5 | -360.55 | <0.001 |
| **Global Efficiency** | 0.340 | 0.006 | 152E5 | 60.50 | <0.001 |
| **Local Efficiency** | 0.161 | 0.004 | 152E5 | 38.33 | <0.001 |
| **Distance** | -0.034 | 0.002 | 152E5 | -19.58 | <0.001 |
| **Distance$^2$** | 0.016 | 0.001 | 152E5 | 11.49 | <0.001 |
| **Age** | -0.016 | 0.009 | 152E5 | -1.73 | 0.084 |
| **Positive Edge Density** | -0.074 | 0.009 | 152E5 | -7.96 | <0.001 |
| **Local Efficiency * Age** | 0.003 | 0.004 | 152E5 | 0.80 | 0.426 |
| **Outgoing Global Efficiency * Age** | -0.016 | 0.006 | 152E5 | -2.90 | 0.004 |

| 4b. Solutions for Fixed Effects – aFN Model 2 (Incoming Connections) | | | | | |
|---|---|---|---|---|---|
| **Effect** | **Estimate** | **Standard Error** | **DF** | **t Value** | **p** |
| **Intercept** | -3.235 | 0.009 | 152E5 | -342.13 | <0.001 |
| **Global Efficiency** | 0.276 | 0.006 | 152E5 | 44.14 | <0.001 |
| **Local Efficiency** | 0.201 | 0.004 | 152E5 | 50.19 | <0.001 |
| **Distance** | -0.038 | 0.002 | 152E5 | -20.48 | <0.001 |
| **Distance$^2$** | 0.015 | 0.001 | 152E5 | 10.47 | <0.001 |
| **Age** | -0.015 | 0.010 | 152E5 | -1.56 | 0.119 |
| **Positive Edge Density** | -0.123 | 0.010 | 152E5 | -12.62 | <0.001 |
| **Local Efficiency * Age** | 0.007 | 0.004 | 152E5 | 1.70 | 0.089 |
| **Incoming Global Efficiency * Age** | -0.014 | 0.006 | 152E5 | -2.20 | 0.028 |

**Supplementary Tables 4a and 4b. Multivariate Mixed Model Results – NCANDA San Diego Site at Baseline - aFN**

Results of multivariate mixed model framework for associating age with connection probability in aFNs. Our model controlled for density in the aFNs. Age had a significant interaction with outgoing global efficiency but not local efficiency.

| 5a. Solutions for Fixed Effects – aFN Model 1 (Outgoing Connections) | | | | | |
|---|---|---|---|---|---|
| **Effect** | **Estimate** | **Standard Error** | **DF** | **t Value** | **p** |
| **Intercept** | -3.218 | 0.008 | 141E5 | -405.38 | <0.001 |
| **Global Efficiency** | 0.325 | 0.006 | 141E5 | 53.45 | <0.001 |
| **Local Efficiency** | 0.184 | 0.004 | 141E5 | 50.24 | <0.001 |
| **Distance** | -0.037 | 0.002 | 141E5 | -19.84 | <0.001 |
| **Distance$^2$** | 0.015 | 0.001 | 141E5 | 10.88 | <0.001 |
| **Age** | -0.007 | 0.008 | 141E5 | -0.92 | 0.357 |
| **Positive Edge Density** | -0.113 | 0.008 | 141E5 | -13.85 | <0.001 |
| **Local Efficiency * Age** | <0.001 | 0.004 | 141E5 | 0.09 | 0.928 |
| **Outgoing Global Efficiency * Age** | -0.011 | 0.006 | 141E5 | -1.82 | 0.069 |

| 5b. Solutions for Fixed Effects – aFN Model 2 (Incoming Connections) | | | | | |
|---|---|---|---|---|---|
| **Effect** | **Estimate** | **Standard Error** | **DF** | **t Value** | **p** |
| **Intercept** | -3.186 | 0.009 | 141E5 | -361.44 | <0.001 |
| **Global Efficiency** | 0.265 | 0.007 | 141E5 | 40.65 | <0.001 |
| **Local Efficiency** | 0.227 | 0.004 | 141E5 | 60.18 | <0.001 |
| **Distance** | -0.043 | 0.002 | 141E5 | -22.30 | <0.001 |
| **Distance$^2$** | 0.014 | 0.001 | 141E5 | 9.38 | <0.001 |
| **Age** | -0.011 | 0.009 | 141E5 | -1.21 | 0.225 |
| **Positive Edge Density** | -0.155 | 0.009 | 141E5 | -16.84 | <0.001 |
| **Local Efficiency * Age** | 0.004 | 0.004 | 141E5 | 1.16 | 0.248 |
| **Incoming Global Efficiency * Age** | -0.020 | 0.007 | 141E5 | -2.99 | 0.003 |

**Supplementary Tables 5a and 5b. Multivariate Mixed Model Results – NCANDA San Diego Site at 1 Year Follow-up - aFN**

Of the 212 participants that completed scans at baseline at the San Diego site, 197 returned for one year follow-up scans. In this sample, age had a near-significant interaction with outgoing global efficiency and a significant interaction with incoming global efficiency.

| 6a. Solutions for Fixed Effects – aFN Model 1 (Outgoing Connections) | | | | | |
|---|---|---|---|---|---|
| **Effect** | **Estimate** | **Standard Error** | **DF** | **t Value** | **p** |
| **Intercept** | -3.195 | 0.010 | 119E5 | -307.06 | <0.001 |
| **Global Efficiency** | 0.323 | 0.006 | 119E5 | 50.33 | <0.001 |
| **Local Efficiency** | 0.200 | 0.004 | 119E5 | 44.66 | <0.001 |
| **Distance** | -0.038 | 0.002 | 119E5 | -17.69 | <0.001 |
| **Distance$^2$** | 0.013 | 0.001 | 119E5 | 9.27 | <0.001 |
| **Age** | -0.008 | 0.010 | 119E5 | -0.76 | 0.448 |
| **Positive Edge Density** | -0.108 | 0.011 | 119E5 | -10.16 | <0.001 |
| **Local Efficiency * Age** | <0.001 | 0.004 | 119E5 | -0.03 | 0.978 |
| **Outgoing Global Efficiency * Age** | -0.013 | 0.006 | 119E5 | -2.00 | 0.046 |

| 6b. Solutions for Fixed Effects – aFN Model 2 (Incoming Connections) | | | | | |
|---|---|---|---|---|---|
| **Effect** | **Estimate** | **Standard Error** | **DF** | **t Value** | **p** |
| **Intercept** | -3.169 | 0.012 | 119E5 | -269.57 | <0.001 |
| **Global Efficiency** | 0.275 | 0.007 | 119E5 | 39.92 | <0.001 |
| **Local Efficiency** | 0.232 | 0.005 | 119E5 | 48.05 | <0.001 |
| **Distance** | -0.042 | 0.002 | 119E5 | -20.35 | <0.001 |
| **Distance$^2$** | 0.013 | 0.002 | 119E5 | 8.31 | <0.001 |
| **Age** | -0.006 | 0.012 | 119E5 | -0.53 | 0.594 |
| **Positive Edge Density** | -0.162 | 0.012 | 119E5 | -13.21 | <0.001 |
| **Local Efficiency * Age** | 0.008 | 0.005 | 119E5 | 1.67 | 0.095 |
| **Incoming Global Efficiency * Age** | -0.009 | 0.007 | 119E5 | -1.36 | 0.173 |

**Supplementary Tables 6a and 6b. Multivariate Mixed Model Results – NCANDA Duke Site at Baseline - aFN**

The Duke University site had 166 participants that completed scans at baseline. In this sample, age had a significant interaction with outgoing global efficiency but not incoming global efficiency.

| Solutions for Fixed Effects - sFN | | | | | |
|---|---|---|---|---|---|
| **Effect** | **Estimate** | **Standard Error** | **DF** | **t Value** | **p** |
| **Intercept** | -3.718 | 0.026 | 758E4 | -141.57 | <0.001 |
| **Global Efficiency** | 1.244 | 0.008 | 758E4 | 151.15 | <0.001 |
| **Local Efficiency** | 0.078 | 0.007 | 758E4 | 10.55 | <0.001 |
| **Distance** | -0.624 | 0.010 | 758E4 | -65.44 | <0.001 |
| **Distance$^2$** | 0.423 | 0.004 | 758E4 | 107.92 | <0.001 |
| **Age** | -0.013 | 0.027 | 758E4 | -0.47 | 0.638 |
| **Positive Edge Density** | -0.047 | 0.027 | 758E4 | -1.75 | 0.081 |
| **Local Efficiency * Age** | -0.020 | 0.007 | 758E4 | -2.67 | 0.008 |
| **Global Efficiency * Age** | -0.008 | 0.008 | 758E4 | -1.01 | 0.313 |

**Supplementary Table 7. Multivariate Mixed Model Results – NCANDA San Diego Site at Baseline - sFN**

Results of multivariate mixed model framework for associating age with connection probability in sFNs. Our model controlled for density in the sFNs. Age had a significant interaction with local efficiency but not global efficiency.
